# Reversal of ATP synthase is a key attribute for cellular differentiation of the *Trypanosoma brucei* insect forms

**DOI:** 10.1101/2025.06.16.659894

**Authors:** Kunzová Michaela, Doleželová Eva, Moos Martin, Panicucci Brian, Zíková Alena

**Affiliations:** Institute of Parasitology, Biology Centre, Czech Academy of Sciences, Ceske Budejovice, Czech Republic; Faculty of Science, University of South Bohemia, Ceske Budejovice, Czech Republic; Institute of Entomology, Biology Centre, Czech Academy of Sciences, Ceske Budejovice, Czech Republic

## Abstract

The mitochondrial F_o_F_1_-ATP synthase is a reversible nanomachine responsible for generating the majority of cellular ATP by oxidative phosphorylation. In various pathological contexts associated with mitochondrial dysfunction, this enzyme can reverse its function to maintain essential mitochondrial membrane potential at the expense of ATP. This reversal is regulated by the conserved protein Inhibitory Factor 1 (IF1). Here, we demonstrate that ATP synthase reversal also occurs under physiological conditions during the cellular differentiation of the unicellular parasite *Trypanosoma brucei*, which transitions between insect and mammalian hosts. Differentiation of the insect forms is marked by upregulation of alternative oxidase (AOX) along with decreased expression of trypanosomal IF1 (TbIF1), collectively creating a metabolic environment conducive to ATP synthase reversal. We show that this reversal is key for proper parasite differentiation: parasites lacking TbIF1 efficiently transitioned into the mammalian-infective form. In these TbIF1 knockout parasites, ATP synthase reversal was associated with an increased ADP/ATP ratio, activation of AMP-activated protein kinase, enhanced cellular respiration driven by elevated proline metabolism, and increased mitochondrial and cellular reactive oxygen species (ROS), known signaling molecules. Conversely, parasites with inducible TbIF1 overexpression failed to reverse ATP synthase activity, showed no AMPK activation or ROS elevation, and remained locked in the initial insect stage. Our findings highlight the critical role of TbIF1 downregulation in enabling life cycle progression and underscore the unique function of the ATP synthase/IF1 axis in cellular signaling.

## Introduction

Mitochondria are multifaceted organelles that play a crucial role in cellular bioenergetics, metabolism, and intracellular signaling [1, 2]. Central to these processes is the F_o_F_1_ ATP synthase (ATP synthase), a molecular nanomachine that synthesizes the majority of cellular ATP via oxidative phosphorylation (OXPHOS) under aerobic conditions [3, 4]. The synthesizing, or forward, mode of this complex is driven by the high proton motive force (pmf) generated by the electron transport chain (ETC) complexes. As with most enzymes, the complex is reversible, and thus can assume a hydrolytic function when the pmf is compromised (e.g., during hypoxia/anoxia conditions, in cells with dysfunctional ETC, or with low electron input) [5, 6]. This hydrolytic activity consumes large amounts of ATP, making it potentially harmful to the cell if not properly regulated. In mitochondria, a unique mechanism involving a short polypeptide named Inhibitory Factor 1 (IF1) has evolved to control the hydrolytic mode of this rotatory machine [7]. The presence of IF1 in almost all aerobic eukaryotes with typical mitochondria suggests that the mechanism regulating ATP synthase activity is conserved among various groups of organisms [8].

The unique characteristic of IF1 is its ability to selectively inhibit the hydrolytic activity of ATP synthase, acting as a unidirectional inhibitor [9]. This inhibition is influenced by environmental conditions such as a pH below 7 and high concentrations of cations (e.g., Ca2+), both of these conditions activate IF1 [10, 11]. The process of IF1 activation involves the exposure of its intrinsically disordered N-terminus, leading to the formation of a short α-helix which then protrudes into the F_1_ moiety of ATP synthase. It forms contacts with the α-helical coiled-coil region of the F_1_-ATPase subunit γ, thereby halting its clockwise rotation (as viewed perpendicular to the mitochondrial membrane plane, with the mitochondrial matrix on top) [12]. When the pmf is restored, mitochondrial membrane potential (ΔΨm) drives the clockwise rotation of the subunit γ, which forcefully ejects IF1 from the F_1_ domain [13]. Notably, it has been proposed that IF1 also suppresses the ATP synthetic activity of the complex in certain human cells [14, 15], although this view has been recently challenged by cell-based and *in vitro* assays and hence remains uncertain [9, 16, 17].

There are many biological roles for IF1 that have been so far described. For example, IF1 plays a key role in inhibiting intermediate assemblies of F_1_ particles during complex assembly [18, 19], or in preserving the cellular ATP pool during ischemic conditions in cardiac tissues [20]. Another significant function of IF1 is its association with the type I dimer of the mammalian ATP synthase complex [21], which can self-assemble into longitudinal rows decorating the rims of lamellar cristae [22, 23]. By binding to ATP synthase and stabilizing the oligomers in these long rows, IF1 levels influence the mitochondrial cristae structure [24–27]. Recently, the pro-oncogenic potential of IF1 has garnered considerable attention, as IF1 expression is elevated in highly proliferative cancer cells and its presence promotes proliferation and resistance to hypoxic conditions and cell death [28]. Other potential mechanisms of IF1 action are relevant to mitophagy, neurodegenerative diseases and aging [16]. Last, but not least, IF1 might be necessary not only under pathophysiological conditions, but also under normal physiological conditions. Accumulating evidence supports the possibility that both ATP synthase activities co-exist within a single coupled intact mitochondrion [29–32]. This underscores the importance of the highly conserved IF1, whose activity might be needed to block ATP hydrolysis under OXPHOS conditions.

In this study, we investigated the role of IF1 under physiological conditions during the extensive metabolic reprogramming underlying cellular differentiation of the unicellular parasite *Trypanosoma brucei* [33]. *T. brucei* belongs to the group of African trypanosomes, which impose significant medical and economic burdens as infectious agents of humans and domestic animals, causing Human and Animal African Trypanosomiasis, respectively [34, 35]. *T. brucei* is a digenetic parasite that alternates between a mammalian host and an insect vector, the tsetse fly. The complex life cycle of these parasites involves a programmed developmental progression through various life cycle stages, each characterized by distinct expression profiles reflecting the parasités specific environments [36, 37].

In the bloodstream of the mammalian host, the bloodstream form of the parasite is covered by a dense variant surface glycoprotein coat [38], and its metabolism is heavily dependent on glucose oxidation via glycolysis [39]. The mitochondrion of this parasite is reduced in size and activity, yet it retains essential functions for example by maintaining the glycolytic redox balance through the dihydroxyacetone/glycerol-3-phosphate shuttle coupled to the reduction of oxygen via an alternative oxidase (AOX) [40]. In addition, the canonical cytochrome-mediated ETC is absent, and the ΔΨm necessary for protein import and ion homeostasis is maintained by ATP synthase operating in reverse [41, 42]. The ATP demands are met by either the supply from the cytosol or the ATP can also be generated in the mitochondrial matrix by substrate phosphorylation [43, 44]. The expression of *T. brucei* IF1 (TbIF1) is fully restricted in the bloodstream forms, and experimentally induced TbIF1 expression is lethal to the parasite [45].

Upon ingestion by the tsetse fly, the bloodstream form parasite rapidly differentiates into the procyclic form, which populates the fly’s midgut. This form is characterized initially by a glycoprotein coat composed of GPEET and EP procyclins, which are subsequently replaced only by EP procyclin [46]. The procyclic form fully utilizes the abundant amino acids, such as proline and threonine, oxidizing them through mitochondrial metabolism via a partial tricarboxylic acid (TCA) cycle [47]. The AOX expression is downregulated, and the majority of electrons released from amino acid oxidation are transferred to oxygen via ETC complexes III and IV, generating the pmf. In this life cycle stage, ATP synthase operates in its canonical mode, synthesizing ATP [48, 49]. In addition to OXPHOS, cellular ATP is also generated by powerful mitochondrial substrate phosphorylation involving the ATP-generating succinyl-Co-A synthetase, which provides the parasite with flexibility in terms of energy metabolism [33, 50]. Procyclic form and bloodstream form parasites are the only two forms commonly grown as proliferating cultures in the laboratory.

Once the infection is fully established in the midgut, the parasite migrates to the salivary glands via proventriculus, transitioning into the epimastigote form, which is characterized by a coat composed of different GPI-anchored proteins called brucei-alanine-rich-proteins (BARPs) [51]. The epimastigote form inhabits the salivary glands and later transforms into the quiescent, cell-cycle arrested metacyclic parasites, which are infectious to the mammalian host. The metacyclic form is pre-adapted for the survival in the mammalian host by adjusted gene expression profile [52, 53]. While the metabolism of the latter two forms remains to be fully elucidated, partial understanding can be derived from studies utilizing *in vitro* differentiation that mimics the developmental processes occurring in the tsetse fly [54, 55]. This differentiation depends on the over-expression of RNA-binding protein 6 (RBP6), whose expression is highly upregulated in salivary gland parasites [56] and, *in vitro*, it triggers the transformation of procyclic parasites into epimastigotes and metacyclics in the process called metacyclogenesis [57, 58]. Omics and cell-based analyses have established that during this differentiation, there is a significant remodeling of the ETC, with AOX expression being strongly upregulated while ETC complexes III and IV are downregulated. Moreover, TbIF1 expression is developmentally downregulated, suggesting a need for ATP synthase reversal during parasite differentiation [55, 59].

Using gain- and loss-of-function experiments, we established that the absence of TbIF1 promotes the differentiation of the parasite to the metacyclic form *in vitro*, while its inducible over-expression halts initial steps of the differentiation. We propose that the absence of TbIF1 allows the reversal of the partial ATP synthase pool promoted by the increased expression of AOX, leading to changes in cellular respiration, reactive oxygen species (ROS) levels, ADP/ATP ratio, and activation of AMP-activated protein kinase (AMPK), an enzyme that acts as an energy sensor in cells. TbIF1 programmed down-regulation is a key adaptive mechanism that enables the ATP synthase reversal under physiological conditions and, as a consequence, TbIF1 modulates *Trypanosoma* energy metabolism during cellular differentiation.

## Methodology

### Cell Lines and Culture Conditions

The RBP6 overexpression (RBP6^OE^) cell line was generated as previously described [55], with expression induced by daily addition of 10 µg/ml tetracycline. To generate the RBP6^OE^_TbIF1dKO line, both alleles of *TbIF1* (Tb927.10.2970) were sequentially disrupted via homologous recombination. For the first allele, 5′ and 3′ UTRs were PCR-amplified from PCF 427 genomic DNA, cloned into the pLew13 vector containing a neomycin resistance cassette and a T7 RNA polymerase gene, linearized with NotI, and transfected into PCF 427 cells using AMAXA nucleofection; positive clones were selected with G418. The second allele was disrupted using pLew90 bearing *TbIF1* intergenic homology arms and a hygromycin resistance cassette with a gene for tetracycline repressor, followed by electroporation into single knockout cells and selection with hygromycin [60]. The RBP6 overexpression construct was then introduced into the double-knockout line. To generate the RBP6^OE^_TbIF1^OE^ cell line, the *TbIF1* coding sequence was PCR-amplified from PCF 427 genomic DNA and cloned into the pLew79 vector for tetracycline-inducible expression. The construct was linearized with NotI, transfected into RBP6^OE^ cells, and puromycin-resistant clones were selected. The PCF 427 and all the generated cell lines were maintained at 27°C in glucose-free SDM-80 medium supplemented with 10% heat-inactivated fetal bovine serum (FBS), 7.5 µg/ml hemin, and 50 mM N-acetyl-D-glucosamine to minimize uptake of residual glucose from FBS. Selection antibiotics were included as appropriate to maintain genetic constructs: G418 (15 µg/ml), hygromycin B (25 µg/ml), puromycin (1 µg/ml), and phleomycin (2.5 µg/ml). The bloodstream form was cultured at 37°C with 5% CO₂ in HMI-11 medium supplemented with 10% FBS.

### Cell Morphology Assessment and Fluorescence Microscopy

For cell morphology analysis and life cycle stages, 5 × 10⁶ cells were harvested by centrifugation at 1,300 × g for 10 minutes at room temperature (RT), washed with 1 ml of 1× phosphate-buffered saline (PBS; pH 7.4), and fixed in 3.7% formaldehyde in PBS. Fixed cells were applied to poly-L-lysine-coated coverslips and incubated for 15 minutes at RT. After three washes with PBS, coverslips were mounted using ProLong Gold Antifade Mountant containing DAPI to visualize nuclear and mitochondrial DNA (kinetoplast). Fluorescence images were acquired using an Axioplan 2 Imaging Universal microscope (Zeiss) equipped with an Olympus DP73 CCD camera. Cell types corresponding to distinct life cycle stages were assigned based on cell size and shape and the relative positioning of the nucleus and kinetoplast. At least 100 cells per time point were scored in a blinded analysis. All experiments were conducted in a minimum of two biological replicates.

### SDS-PAGE and Western Blotting

Protein samples from whole-cell lysates (1 × 10^7^ cells per lane) were separated by SDS-PAGE and transferred to polyvinylidene difluoride (PVDF) membranes. Membranes were probed with appropriate monoclonal or polyclonal primary antibodies, followed by horseradish peroxidase (HRP)-conjugated anti-mouse or anti-rabbit secondary antibodies. Protein bands were visualized using the Pierce enhanced chemiluminescence (ECL) detection system, and signals were captured using a ChemiDoc imaging system (Bio-Rad). Protein sizes were determined by comparison to the PageRuler prestained protein ladder. A commercially available anti–phospho-AMPKα1/2 (Thr172) polyclonal antibody (Sigma-Aldrich) was used at a 1:1000 dilution. All other antibodies were either previously acquired or generated using His-tagged recombinant proteins, with final production performed by Davids Biotechnologie (Regensburg, Germany). Primary antibodies used in this study included: mouse monoclonal anti–mitochondrial HSP70 (1:5000) [61], rabbit polyclonal anti-RBP6 (1:1000, this work), anti-GPEET (1:1000; a generous gift from Prof. Isabel Roditi), anti-BARP (1:1000, a generous gift from Prof. Isabel Roditi), anti-RBP10 (1:1000; this work), mouse monoclonal anti-AOX (1:500; a generous gift from Prof. Minu Chaudhuri), and rabbit polyclonal antibodies against Rieske (1:1000 [55]), trCOIV (1:1000 [55]), ATP synthase subunit β (1:2000 [62]), p18 (1:1000 [62]), TbIF1 (1:100 [45]), and NDUFA6 (1:1000 [55]).

### ADP/ATP ratio

The cellular ADP/ATP ratio was measured using a luciferase-based enzymatic assay kit (Sigma-Aldrich) to assess the energetic status of the cells. For each experiment, 5 × 10⁶ cells were harvested by centrifugation at 1,500 × g for 10 minutes at RT, then resuspended in 1 ml of 1× PBS; pH 7.4. A total of 1 × 10⁶ cells per well were transferred into a white 96-well plate, and the assay was performed according to the manufacturer’s instructions. Luminescence was measured using a Tecan Infinite M200 plate reader. Each condition was analyzed in both technical and biological triplicates.

### ATP analysis by LC-HRMS

Cells (5×10⁷) were harvested (1300g, 10 min, 4°C), the supernatant removed, and the pellet resuspended in 80 μL cold MeOH:ACN:Water (2:2:1, v/v/v). After 10 min in a 0°C ultrasonic bath, the mixture was centrifuged (7000 rpm, 10 min, 4°C). The supernatant was diluted 1:10 with 50% ACN. ^13C₁₀-ATP (20 ng/sample) was added as internal standard (IS) to LC/MS vials, dried under nitrogen, and reconstituted in 40 μL of the diluted extract for LC-HRMS analysis; remaining extract was stored at –80°C.

ATP quantification followed a previously described liquid chromatography high resolution mass spectrometry (LC-HRMS) method [63] using an Orbitrap QExactive Plus with a Dionex Ultimate 3000 and open autosampler (Thermo Fisher Scientific). The QExactive operated in negative ESI mode (full MS scan, 70–1000 Da) at 70,000 resolution, 3×10⁶ AGC, and 100 ms injection time. Ionization settings included ±3000 V spray voltage, 350°C capillary/probe temperatures, sheath/auxiliary/spare gases at 60/20/1 au, and S-lens at 60 au. Chromatographic separation was performed on a SeQuant ZIC-pHILIC column (150×4.6 mm, 5 μm, Merck) at 35°C, 450 μL/min, 5 μL injection. Mobile phase: acetonitrile (A) and 20 mmol/L ammonium carbonate (B, pH 9.2), with gradient: 0 min, 20% B; 20 min, 80% B; 20.1 min, 95% B; 23.3 min, 95% B; 23.4 min, 20% B; 30 min, 20% B. Data were acquired using Xcalibur v4.0. Each condition was analyzed at least in biological triplicates.

### Measurement of Cellular and Mitochondrial ROS and Mitochondrial Membrane Potential (Δ𝚿m)

Cellular and mitochondrial ROS levels were assessed using carboxy-2′,7′-dichlorofluorescein diacetate (DCF-DA) and MitoSOX Red, respectively. A total of 1 × 10⁷ cells were incubated under standard cultivation conditions (27°C, shaking) with 10 µM DCF-DA (Sigma-Aldrich) for detection of general cellular ROS, or with 5 µM MitoSOX Red Mitochondrial Superoxide Indicator (Thermo Fisher Scientific) for detection of mitochondrial superoxide. After 30 minutes incubation, cells were harvested by centrifugation at 1,500 × g for 10 minutes at RT, washed once with 1 ml of 1× PBS; pH 7.4, and resuspended in 2 ml of PBS. Fluorescence from 10,000 events per sample was measured immediately using a BD FACS Canto II flow cytometer (BD Biosciences).

To assess ΔΨm , cells were stained with 60 nM tetramethylrhodamine ethyl ester (TMRE; Thermo Fisher Scientific) under cultivation conditions for 30 minutes. As a control for mitochondrial depolarization, cells were treated with 20 µM of the protonophore carbonyl cyanide-p-trifluoromethoxyphenylhydrazone (FCCP). Flow cytometry data were analyzed using FlowJo software version 10 (BD Biosciences).

### Measurement of Oxygen Consumption by High-Resolution Respirometry

Cellular oxygen consumption was measured using an O2k high-resolution respirometer (Oroboros Instruments) in MiRO5 respiration medium at 27°C. Each chamber was loaded with 2 × 10⁷ cells. Mitochondrial respiration was stimulated by the addition of either 5 mM proline or 10 mM glycerol-3-phosphate. To differentiate between cytochrome c oxidase (complex IV)–mediated and alternative oxidase (AOX)–mediated respiration, cells were sequentially treated with 1 mM potassium cyanide (KCN) and 250 µM salicylhydroxamic acid (SHAM), respectively.

### *In Vitro* Differentiation of Metacyclic Form to Bloodstream Form

To achieve full differentiation from *T. brucei* metacyclic to bloodstream-form, a rodent-free in vitro protocol adapted from cyclical transmission models was used [64]. Briefly, a four-day tetracycline-induced RBP6^OE^ and RBP6^OE^_TbIF1dKO cultures containing metacyclic forms was centrifuged at 1,300 × g for 10 minutes at RT and resuspended at a density of 5 × 10⁵ cells/ml in differentiation medium composed of Minimal Essential Medium with Earle’s salts, supplemented with 1% non-essential amino acids, 2 g/l glucose, and 15% heat-inactivated rabbit serum (Sigma-Aldrich). Cells were incubated in this medium for 3 hours at 37°C in a closed-lid environment. Following incubation, cells were pelleted again (1,300 × g, 10 minutes, RT) and resuspended in HMI-11 medium supplemented with 1.1% methylcellulose and 10% FBS. Cultures were maintained at 37°C with 5% CO₂ for up to two weeks. Upon the emergence of dividing long slender bloodstream forms, the culture was diluted into HMI-11 medium containing 10% FBS.

### Blue Native PAGE and Immunoblotting of F_o_F_1_ ATP synthase

Blue native polyacrylamide gel electrophoresis (BN-PAGE) followed by western blotting was performed to separate protein complexes in their native conformation, using a protocol adapted from [48]. Mitochondria were isolated from 3 × 10⁸ cells by hypotonic lysis and resuspended in solubilization buffer containing 750 mM aminocaproic acid, 50 mM Bis-Tris, 0.5 mM EDTA (pH 7.0), supplemented with a complete EDTA-free protease inhibitor cocktail and 2% n-dodecyl-β-D-maltoside (DDM). The mixture was incubated on ice for 1 hour to ensure membrane solubilization. Following centrifugation at 16,000 × g for 30 minutes at 4°C, the supernatant was collected and protein concentration was determined using the BCA assay. A total of 8 µg of protein was mixed with a Coomassie-based loading buffer (50 mM aminocaproic acid, 0.5% [w/v] Coomassie Brilliant Blue G-250) and loaded onto 3–12% Bis-Tris Native PAGE gels (Thermo Fisher Scientific). Electrophoresis was carried out at 150 V for 3 hours at 4°C. Proteins were transferred to PVDF membranes and probed with specific antibodies as described above.

### Transmission Electron Microscopy for Mitochondrial Ultrastructure Analysis

Total of 1 × 10⁶ cells were harvested by centrifugation at 620 × g for 10 minutes at RT and fixed in 2.5% glutaraldehyde in 0.1 M phosphate buffer (pH 7.2). Post-fixation was carried out with 2% osmium tetroxide for 2 hours at 4°C. Samples were then washed and dehydrated through a graded acetone series and embedded in Polybed 812 resin (Polysciences, Inc.). Ultrathin sections were prepared using a Leica UCT ultramicrotome (Leica Microsystems), and images were acquired with Transmission Electron Microscope JEM 1400 Flash (JEOL) equipped with a XAROSA camera (SIS).

## Results

### Levels of TbIF1 regulates the RBP6 induced progression through *T. brucei* development

Given that the IF1 is a master regulator of mitochondrial physiology and that levels of TbIF1 expression are downregulated during the development in the tsetse fly [37] as well as following the RBP6 induction (Figure 1A, left panel) [55], we were interested in its role in the parasite differentiation. To this end, we generated procyclic cell lines in which TbIF1 was either lacking or inducible expressed at levels above its endogenous expression (Figure 1A). To generate a cell line lacking TbIF1, we proceeded to remove both TbIF1 alleles by homologous recombination. This was followed by the verification of the successful replacement of both alleles with cassettes containing T7 RNAP and TetR by PCR (Supplementary Figure S1). Subsequently, a cassette containing the RBP6 gene under the control of T7 RNAP and TetR was introduced to allow for the tetracycline inducible expression of RBP6 (RBP6^OE^_TbIF1dKO). To generate a cell line with high expression of TbIF1 during RBP6-induced differentiation, a cassette ensuring tetracycline inducible overexpression of TbIF1 was transfected into the RBP6^OE^ cell line (RBP6^OE^_TbIF1^OE^). The RBP6-driven differentiation of the generated cell lines was induced by tetracycline. The induced RBP6 and TbIF1 expression was verified by Western blot. As anticipated, TbIF1 expression was undetectable in RBP6^OE^_TbIF1dKO, while it remained consistently elevated in RBP6^OE^_TbIF1^OE^ due to the constant presence of tetracycline in the medium (Figure 1A, middle and right panels).

**Figure 1.**
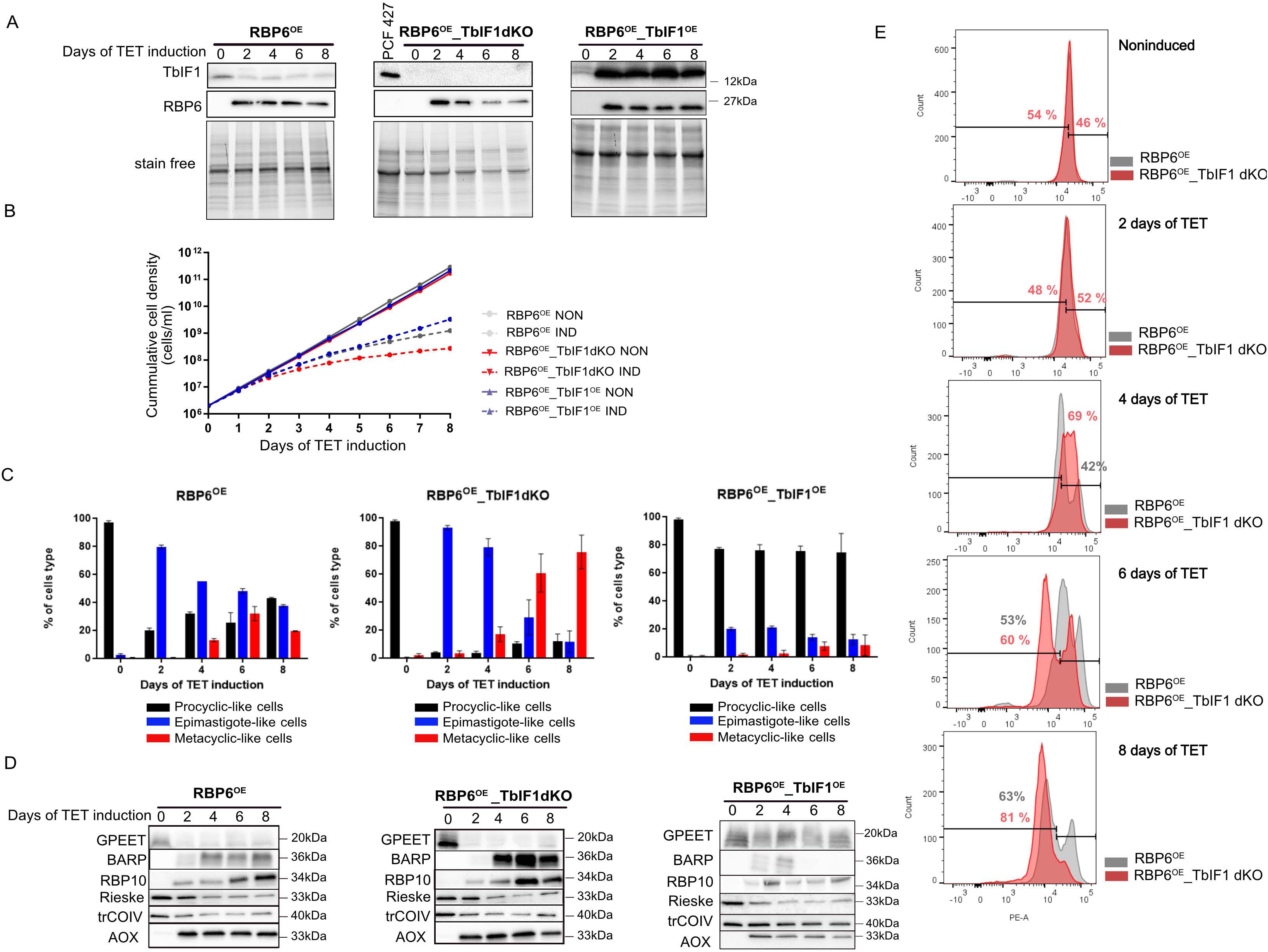
Lack of TbIF1 enhances differentiation of procyclic *T. brucei* cells. (A) Western blot analysis of whole cell lysates from the parental RBP6^OE^, RBP6^OE^_TbIF1dKO and RBP6^OE^_TbIF1^OE^ cells harvested at timepoints 0, 2, 4, 6 and 8 after the tetracycline (TET)-induced RBP6 overexpression. The procyclic strain PCF 427 we used as a control for IF1 detection. Stain free gels served as a loading control for equal loading of 1 x 10^7^ cells/well. (B) Growth curves of RBP6^OE^, RBP6^OE^_TbIF1dKO and RBP6^OE^_TbIF1^OE^ measured for 8 days. (C) Cell type scoring analysis for the presence procyclic-like of epimastigote-like and metacyclic-like cells upon induction of RBP6 overexpression in all three cell lines. (D) Western blot analysis of whole cell lysates from RBP6^OE^, RBP6^OE^_TbIF1dKO and RBP6^OE^_TbIF1^OE^ cells undergoing differentiation using a panel of various antibodies. (E) Flow cytometry overlay histograms for TMRE-stained cell population of RBP6^OE^ and RBP6^OE^_TbIF1dKO noninduced and induced for 2, 4, 6 and 8 days.

A typical manifestation of RBP6-induced differentiation is a slowdown in cell growth, as the parasite differentiates from dividing procyclic form and epimastigotes to the metacyclic form that is arrested in the G1/G0 cell cycle phase [58]. The RBP6^OE^_TbIF1dKO line exhibited a more pronounced growth defect compared to the RBP6^OE^ cell line, whereas the RBP6^OE^_TbIF1^OE^ cell line showed the least severe growth defect (Figure 1B). To gain insight into the dynamics of the differentiation process, we determined the individual life cycle forms (procyclic, epimastigote and metacyclic). We took into considerations stage-specific morphology cues: such as shape and cell size as well as the relative position of the kinetoplast to the nucleus. Procyclic cells have their mitochondrial DNA (kDNA) located in the posterior part of the cell. During the maturation of epimastigotes, their kDNA migrates and it is typically located near the nucleus at the anterior part of the cell. In metacyclic trypomastigotes, which are smaller than both procyclics and epimastigotes, the kinetoplast is located at the more rounded posterior tip (Supplementary Figure S2). The RBP6^OE^ showed a typical differentiation profile [55] with almost 80% of epimastigotes on day 2 and with close to 40% of metacyclic parasites at day 6. In accordance with the growth curves, RBP6^OE^_TbIF1dKO demonstrated enhanced differentiation efficiency, with the detection of 60-80% metacyclic trypomastigotes at days 6 and 8, which was significantly higher than what observed in RBP6^OE^. In contrast, RBP6^OE^_TbIF1^OE^ exhibited a significant impairment of differentiation to epimastigotes, with less than 20% of the culture reaching this stage at day 2 post induction (Figure 1C).

RBP6-induced differentiation is accompanied by a change in the expression profile, with some marker proteins being readily visualised using available antibodies. In RBP6^OE^, the expression of the surface glycoprotein GPEET is downregulated and later replaced by BARP, which signals the presence of mature epimastigotes. Additionally, the expression of RNA binding protein 10 (RBP10), a protein whose expression correlates with that of the bloodstream-form like gene profile [65], is increased during differentiation. As a consequence of the remodeling of the canonical ETC, the subunits of complexes III and IV, Rieske and trCOIV, respectively, are downregulated, while the subunit of the AOX is strongly upregulated shortly after the induction of RBP6 overexpression (Figure 1D, left panel). In the case of RBP6^OE^_TbIF1dKO, a more pronounced increase in BARP and RBP10 signals was observed, which is consistent with the growth curves indicating a higher proportion of mature epimastigotes and metacyclics in the culture (Figure 1D, middle panel). In contrast, for RBP6^OE^_TbIF1^OE^, there was no decrease in GPEET and the signal for BARP was only barely discernible on days 2 and 4 following the induction of RBP6^OE^. The remaining markers exhibited a comparable trend to that observed in RBP6^OE^, albeit to a lesser extent. This suggests that RBP6-driven programmed differentiation was triggered on the molecular level in the RBP6^OE^_TbIF1^OE^ cells. However, the transition into the next life cycle stage was prohibited by the TbIF1 overexpression as the procyclic cells appear to be the dominant cell population in the culture (Figure 1D, right panel).

Furthermore, the enhanced differentiation of RBP6^OE^_TbIF1dKO is also corroborated by the histograms of TMRE-stained cells, which are employed for the estimation of ΔΨm. Figure 1E illustrates the typical pattern of RBP6^OE^ cells (represented by grey peaks) when a distinct cell population exhibits increased TMRE fluorescence intensity on days 4 and 6. Inversely, another distinct cell population exhibits a decrease in ΔΨm on day 8 (63 ± 7%). In RBP6^OE^_TbIF1dKO, this pattern is even more pronounced on day 4, 69 ± 6 % of cells exhibit increased TMRE fluorescence, while on day 8, 81 ± 8 % of cells are present in the population with lower ΔΨm. When these results are combined with cell type scoring (Figure 1C) it indicates that BARP-expressing epimastigotes exhibit a higher ΔΨm, whereas metacyclics arrested in the cell cycle exhibit a lower ΔΨm.

These findings indicate that during RBP6-driven differentiation, TbIF1 plays a pivotal role in the process of programmed development. The absence of TbIF1 is supporting metacyclogenesis, whereas the presence of TbIF1 significantly impairs the parasite’s capacity to differentiate into epimastigotes let alone infectious metacyclic form. Given that the primary function of IF1 is to inhibit the reversal of ATP synthase, this activity appears to be an essential attribute during parasite differentiation.

### TbIF1 modulates changes in mitochondrial functions during parasite differentiation

To gain insight into the mechanism by which the absence of TbIF1 allows procyclic cells to progress more efficiently through differentiation and to ascertain the role of ATP synthase in this process, we examined typical mitochondrial functions (i.e., complexes III/IV- and AOX-mediated respiration, mitochondrial reactive oxygen species (mROS) levels, and ΔΨm) in the RBP6^OE^_TbIF1dKO cell line in comparison with RBP6^OE^. One of the earliest hallmarks of RBP6-induced differentiation is the increased expression of AOX, which is observed as early as 12 hours after the tetracycline induction (Figure 2A). Generally, AOX competes with complex III for electrons from ubiquinol, subsequently transferring them directly to oxygen; although, it does not contribute to the pmf. The elevated levels of AOX were accompanied by increase in proline-based respiration in live intact cells on day 2 in both cell lines, suggesting that this amino acid is being consumed and oxidized at a higher rate, leading to an increase in the electron flow to the ETC. The surplus of electrons entering the ETC were passed to AOX (Figure 2B). It is notable that the RBP6^OE^_TbIF1dKO exhibited a markedly elevated oxygen consumption rate on day 2 in comparison to RBP6^OE^ (231 ± 11 vs. 124 ± 12 pmol/(s*ml), respectively). This was despite the levels of AOX remained comparable between these two cell lines (Figure 2C) suggesting that the loss of TbIF1 may be a contributing factor to this effect. The elevated mitochondrial respiration did not result in discernible changes in ΔΨm at day 2 of RBP6 induction in either cell line as measured in a cell population stained with TMRE (Figure 2D), but was accompanied by augmented production of mROS, predominantly superoxide, as evidenced by the MitoSOX dye. The RBP6^OE^_TbIF1dKO exhibited heightened levels of MitoSOX fluorescence, which aligns with the observed increase in oxygen consumption (Figure 2E).

**Figure 2.**
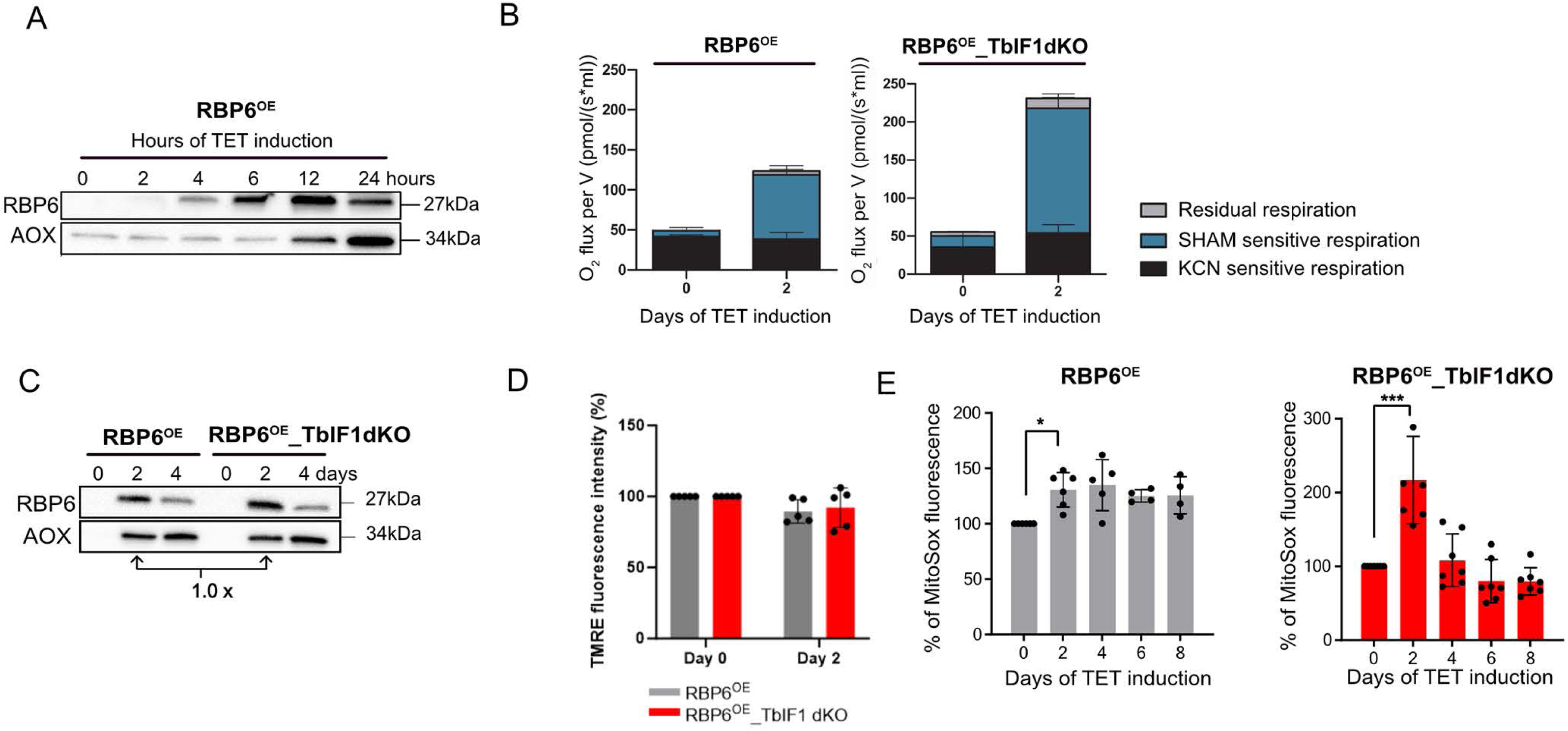
Absence of TbIF1 during *T. brucei* procyclic differentiation enhances oxygen consumption and mROS generation. (A) Western blot analysis of whole cell lysates from RBP6^OE^ cells induced for 0, 2, 4, 6, 12 and 24 hours. (B) Oxygen consumption rates in the presence of 5 mM proline in intact RBP6^OE^ and RBP6^OE^_TbIF1dKO cells induced for 0 and 2 days. The ratio between complex IV- and AOX-mediated respiration was determined using KCN, a potent inhibitor of complex IV, and SHAM, a potent inhibitor of AOX. (means ± s.d., n = 3) (C) Western blot analysis of whole-cell lysates from RBP6^OE^ and RBP6^OE^_TbIF1dKO induced for 0, 2 and 4 days using antibodies against RBP6 and AOX. (D) Flow cytometry analysis of TMRE-stained RBP6^OE^ and RBP6^OE^_TbIF1dKO cells induced for 2 days. (means ± s.d., n= 5) (E) Flow cytometry analysis of MitoSox treated cells to detect mROS levels. Individual values shown as dots. (means ± s.d., n= 4-7, * *P* < 0.05, *** *P* < 0.001)

In mammalian cells, IF1 plays a role in the stability of ATP synthase type I dimer and oligomer, hence, influencing the cristae ultrastructure. Therefore, any alterations in mitochondrial bioenergetics observed in its absence can be attributed to the loss of ATP synthase dimers/oligomers and the subsequent impact on the cristae ultrastructure. To determine whether TbIF1 plays a role in maintaining the stability of ATP synthase type IV dimer in *T. brucei*, we conducted a steady-state analysis of ATP synthase dimers using Blue Native (BN) electrophoresis in RBP6^OE^, RBP6^OE^_TbIF1dKO, and RBP6^OE^_TbIF1^OE^. Our findings indicated that there were no discernible differences in the stability of the ATP synthase dimers or any obvious changes in cristae shapes (Supplementary Figure S3). Therefore, it can be concluded that the effect of increased oxygen consumption and mROS levels is likely due to the reversal of the ATP synthase, which is enabled by a decrease (in tetracycline-induced RBP6^OE^) or absence of TbIF1 (RBP6^OE^_TbIF1dKO). Additionally, the elevated expression of AOX induces a local depolarization of the mitochondrial inner membrane, which may facilitate the reversal of the ATP synthase.

### Absence of TbIF1 allows reversal of ATP synthase

We sought to investigate the extent of ATP synthase reversal in RBP6^OE^ and RBP6^OE^_TbIF1dKO cells at the 0 and 2 days after RBP6 induction. To this end, an *in vitro* assay employing digitonin-permeabilized cells and the dye Safranine O was utilized. The Safranine O fluorescence quenching is an indicator of ΔΨm establishment. NADH-producing substrates (α-ketoglutarate/malate) were added in order to create conditions under which electrons could enter the ETC via alternative dehydrogenase, complex I or complex II, and generate ΔΨm via complexes III/IV that is KCN sensitive. Then ATP was added to facilitate ATP synthase reversal. In RBP6^OE^ noninduced cells, α-ketoglutarate/malate resulted in a certain degree of fluorescence quenching of safranine O, and the addition of ATP led to a further increase in ΔΨm. A proportion of this membrane polarization was oligomycin-sensitive, indicating that ATP synthase reversal may be a contributing factor (Figure 3A). At day 2 after RBP6 induction, when AOX expression is significantly increased (Figure 2B), the NADH-producing substrates induced lower polarization, suggesting that electrons are partially diverted to non-proton-pumping AOX (Figure 3B). In RBP6^OE^_TbIFdKO noninduced cells, both substrates induced similar polarization as in RBP6^OE^ noninduced, but in the absence of TbIF1, the proportion of polarization generated by ATP synthase was slightly greater (Figure 3C and E). Finally, in RBP6^OE^_TbIF1dKO induced for 2 days, the NADH-producing substrates induced similar low polarization as in RBP6^OE^ induced cells, but the ATP-induced polarization was significantly greater and almost fully abolished by oligomycin (Figure 3D and E).

**Figure 3.**
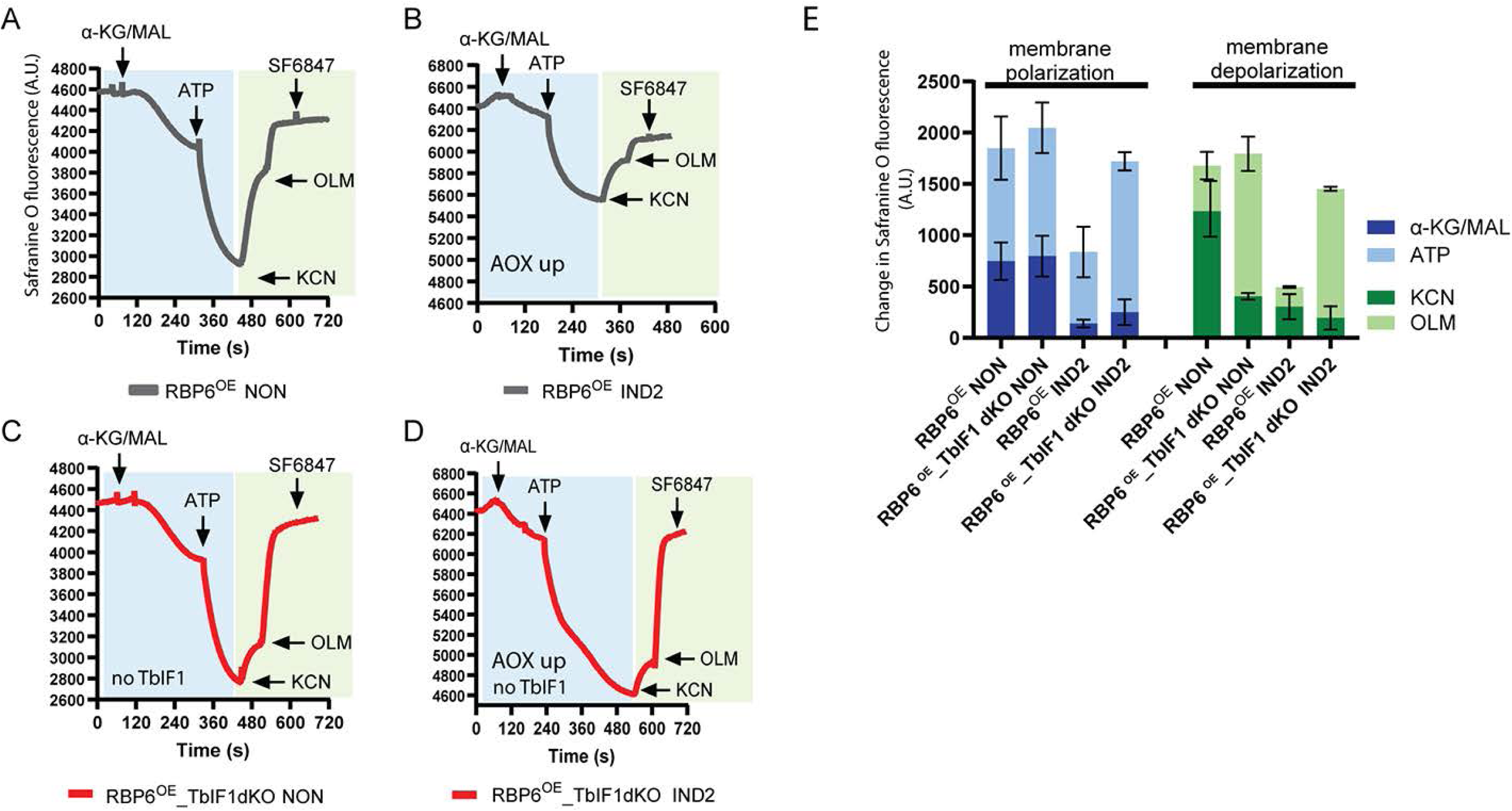
Reversal of *T. brucei* ATP synthase in the absence of TbIF1. (A-D) Representative traces of *in situ* generation and dissipation of the ΔΨm in response to substrates and inhibitors in RBP6^OE^ and RBP6^OE^_TbIF1dKO cells induced for 0 and 2 days. Substrates: α-KG/malate (α-KG/MAL), ATP, ADP; inhibitors: KCN, oligomycin (OLM) and protonophore SF6847. (E) Changes in Safranine O fluorescence after the addition of α-KG/MAL and ATP (membrane polarization, blue columns) and KCN and OLM (membrane depolarization, green columns). (means ± s.d., n=3)

The results show that in the presence of the external amount of ATP, the ATP synthase is able to reverse and contribute to the ΔΨm, in addition to the canonical ETC. This phenomenon is even more pronounced in the presence of AOX, which causes lower membrane polarization, and in the absence of TbIF1, which allows full ATP synthase reversal. This is consistent with recently published data showing that there are distinct populations of ATP synthase in mitochondria that can function either synthetically or hydrolytically, depending on the local pmf [66]. The reciprocal relationship between these activities then determines the bioenergetic outcome of the whole mitochondrion.

### The reversed activity of ATP synthase contributes to the ΔΨm during the differentiation

Our results demonstrate that ATP synthase in digitonin-permeabilized parasites is capable of efficient reversal and contributes to ΔΨm under conditions of high AOX expression and ATP excess. It would be of interest to determine whether this phenomenon occurs under physiological conditions in living cells undergoing differentiation. To address this question, differentiation was triggered by tetracycline in RBP6^OE^_TbIF1^OE^ cell lines, in which the increased expression of TbIF1 is also triggered (Figure 1A). The ΔΨm was measured in the cell population by flow cytometry using the TMRE dye. The results demonstrated a reduction in TMRE fluorescence on the second day following RBP6 induction (Figure 4A-B). Since AOX is detectable in the Western blot on this day (Figure 1D, right panel), it can be concluded that the presence of TbIF1 does not allow the reversal of ATP synthase to partially restore the ΔΨm dissipated by the presence of AOX. The RBP6 induction resulted in slightly higher oxygen consumption (Figure 4C) and higher mitochondrial ROS levels (Figure 4D), but to a much lesser extent, suggesting that the presence of TbIF1 interferes with the RBP6-induced remodeling.

**Figure 4.**
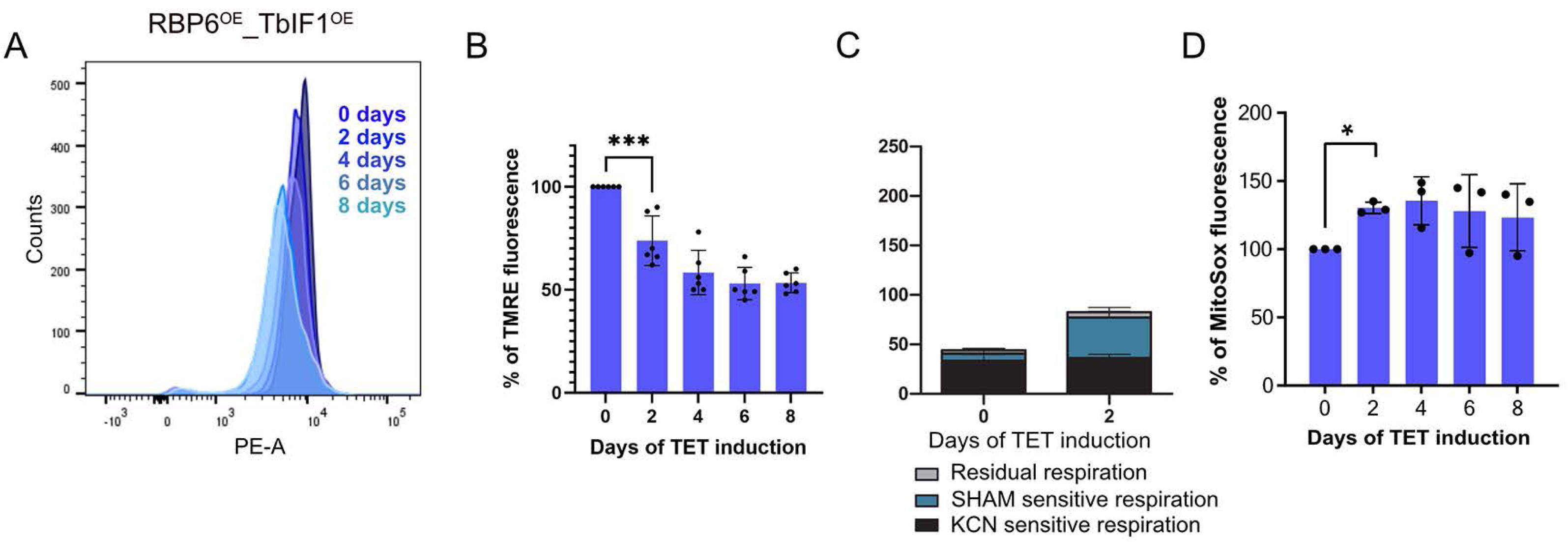
TbIF1 overexpression in RBP6OE cells leads to lowered. ΔΨm. (A) A representative flow cytometry histogram of TMRE-stained RBP6^OE^_TbIF1^OE^ cells induced for 0, 2, 4, 6 and 8 days. (B) Flow cytometry analysis of TMRE-stained RBP6^OE^_TbIF1^OE^ cells induced for 0, 2, 4, 6 and 8 days. Individual values shown as dots. (means ± s.d., n= 5, *** *P* < 0.001) (C) Oxygen consumption rates in the presence of 5 mM proline in intact RBP6^OE^_TbIF1^OE^ cells induced for 0 and 2 days. The ratio of complex IV- and AOX-mediated respiration was determined using KCN, a potent inhibitor of complex IV, and SHAM, a potent inhibitor of AOX. (n= 3) (D) Flow cytometry analysis of MitoSox treated cells to detect mROS levels. Individual values shown as dots. (means ± s.d., n= 3, \**P* < 0.05)

Taken together, our data indicate that the programmed downregulation of TbIF1 expression during the RBP6-induced differentiation serves to facilitate the reverse mode of ATP synthase, which enables energetic adaptation and effectively supports parasite differentiation.

### AMPK is activated during the RBP6 OE differentiation

Having demonstrated that the presence of AOX and the absence of TbIF1 facilitate partial reversal of ATP synthase to maintain ΔΨm upon ATP consumption, we sought to further investigate whether the cellular ATP levels and ADP/ATP ratio are altered. The ADP/ATP ratio is monitored by the cell in a sensitive manner, as an elevated value indicates an energy crisis that leads to the activation of a number of signaling pathways, including the activation of AMP activated kinase (AMPK). Indeed, while total ATP levels remained unaltered, the ADP/ATP ratio exhibited a statistically significant elevation from day 1 following RBP6 induction in the RBP6^OE^_TbIF1dKO (Figure 5A-B). The use of an anti-phospho-Thr-172 antibody, which recognizes both TbAMPK α subunits, enabled the detection of AMPK subunit α1 phosphorylation, an indicator of AMPK activation [67], in both cell lines, albeit at an earlier time point in RBP6^OE^_TbIF1dKO. In contrast, no evidence of AMPK subunit α1 activation was observed in RBP6^OE^_TbIF1^OE^ cells that did not undergo differentiation into metacyclic cells (Figure 5C). These findings suggest that AMPK plays a key role in the differentiation of the parasite into quiescent metacyclic parasites.

**Figure 5.**
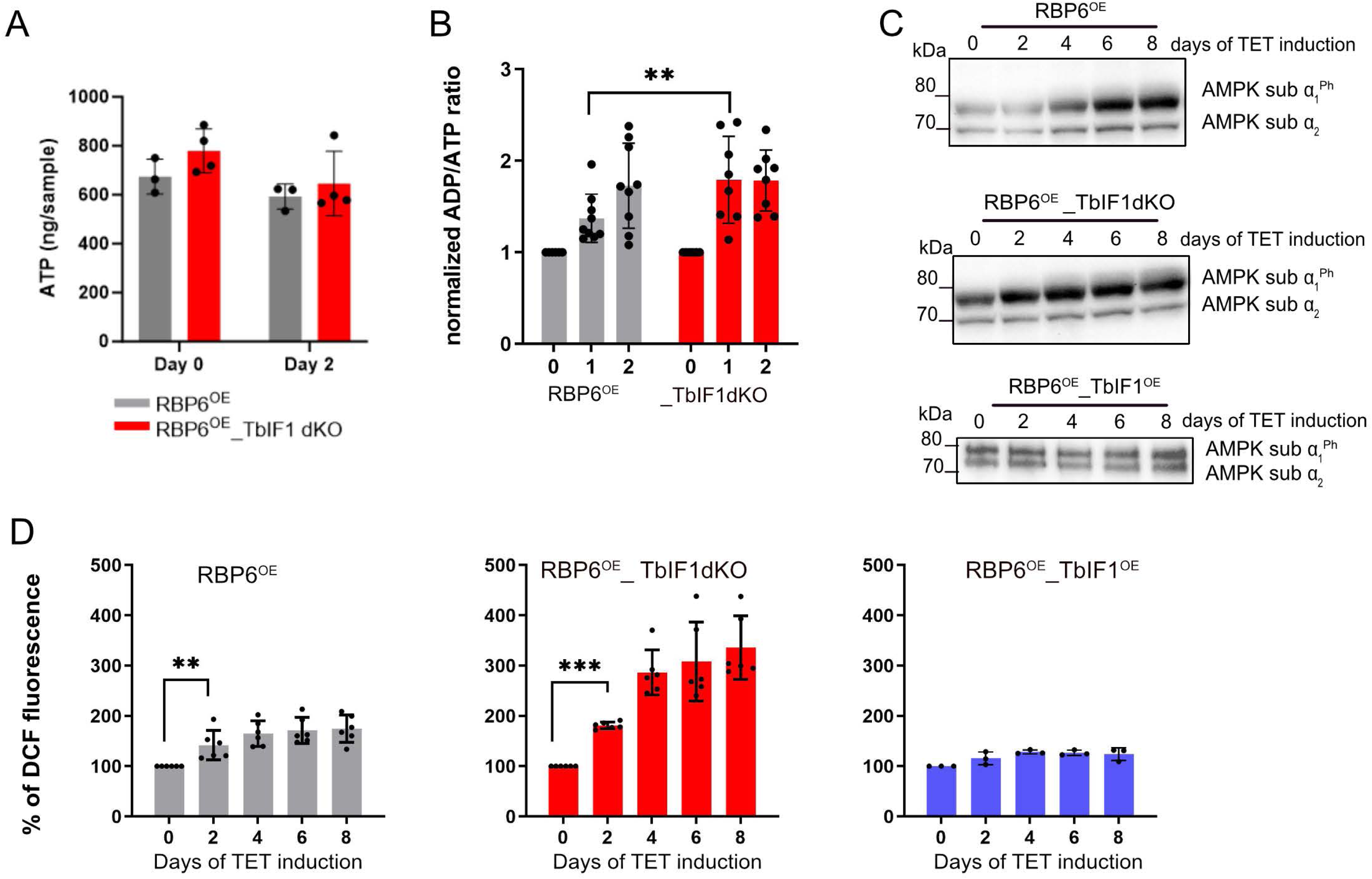
AMPK is activated during *T. brucei* differentiation. (A) Steady-state levels of cellular ATP determined by mass spectrometry in intact RBP6OE and RBP6OE_TbIF1dKO cells. (means ± s.d., n= 3-4) (B) Relative ADP/ATP ratios determined in RBP6^OE^ and RBP6OE_TbIF1dKO expressed as fold increase (means ± s.d., n= 6-9, \*\**P* < 0.01) (C) Western blot analysis of whole-cell lysates from RBP6^OE^ and RBP6OE_TbIF1dKO cells induced for 0, 2, 4, 6 and 8 days using a commercially available anti-phospho-Thr172 AMPK antibody recognizing both AMPK activated α1 and α2 subunits. Signal for AMPK α2 serves as a loading control. (D) Flow cytometry analysis of carboxy-DCF-DA treated cells to detect cellular ROS levels. Individual values shown as dots. (means ± s.d., n= 6, \*\**P* < 0.01, \*\*\**P* < 0.001)

In addition to alterations in the ADP/ATP ratio, AMPK kinase can also be triggered by elevated levels of ROS within cells [68, 69], which act in this case as signaling molecules. It is notable that, in contrast to the augmented cellular ROS levels observed in RBP6^OE^ cells, a considerably greater elevation was detected in RBP6^OE^_TbIF1dKO cells, which progress through differentiation into metacyclics more efficiently. In contrast, no increase in cellular ROS levels was observed in RBP6^OE^_TbIF^OE^ cells, whose differentiation is significantly impaired (Figure 5D).

### The absence of TbIF1 allows the differentiation from metacyclic cells into the long slender bloodstream form

During the differentiation process, we observed a progressive silencing of TbIF1 expression, whose absence is necessary for the initiation and maintenance of the infection in the mammalian host. This is because the bloodstream form stages only utilize the hydrolytic activity of the ATP synthase and TbIF1 expression is lethal to them [45].

In order to fully close the *T. brucei* cycle *in vitro*, we attempted to differentiate metacyclics obtained by RBP6 induction from the parental line (RBP6^OE^) and from the RBP6^OE^_TbIF1dKO cell line *in vitro*. The purified metacyclic form parasites were placed in differentiation medium [64] and subsequently transferred to the HMI-11 medium developed for bloodstream form [70]. Despite repeated attempts, the RBP6^OE^-generated metacyclics were unable to yield viable culture (Figure 6A). Nevertheless, the metacyclics derived from the RBP6^OE^_TbIF1dKO consistently differentiated to viable bloodstream form culture that respires solely through AOX and has downregulated ETC complexes III and IV, as evidenced by western blot analysis (Figure 6 B, C). The capacity to differentiate from metacyclics to BSF *in vitro* provides the opportunity to study the remodeling of the parasite’s cellular metabolism and ultrastructure upon the entry into the mammalian environment, a part of the life cycle that has not been previously investigated.

**Figure 6.**
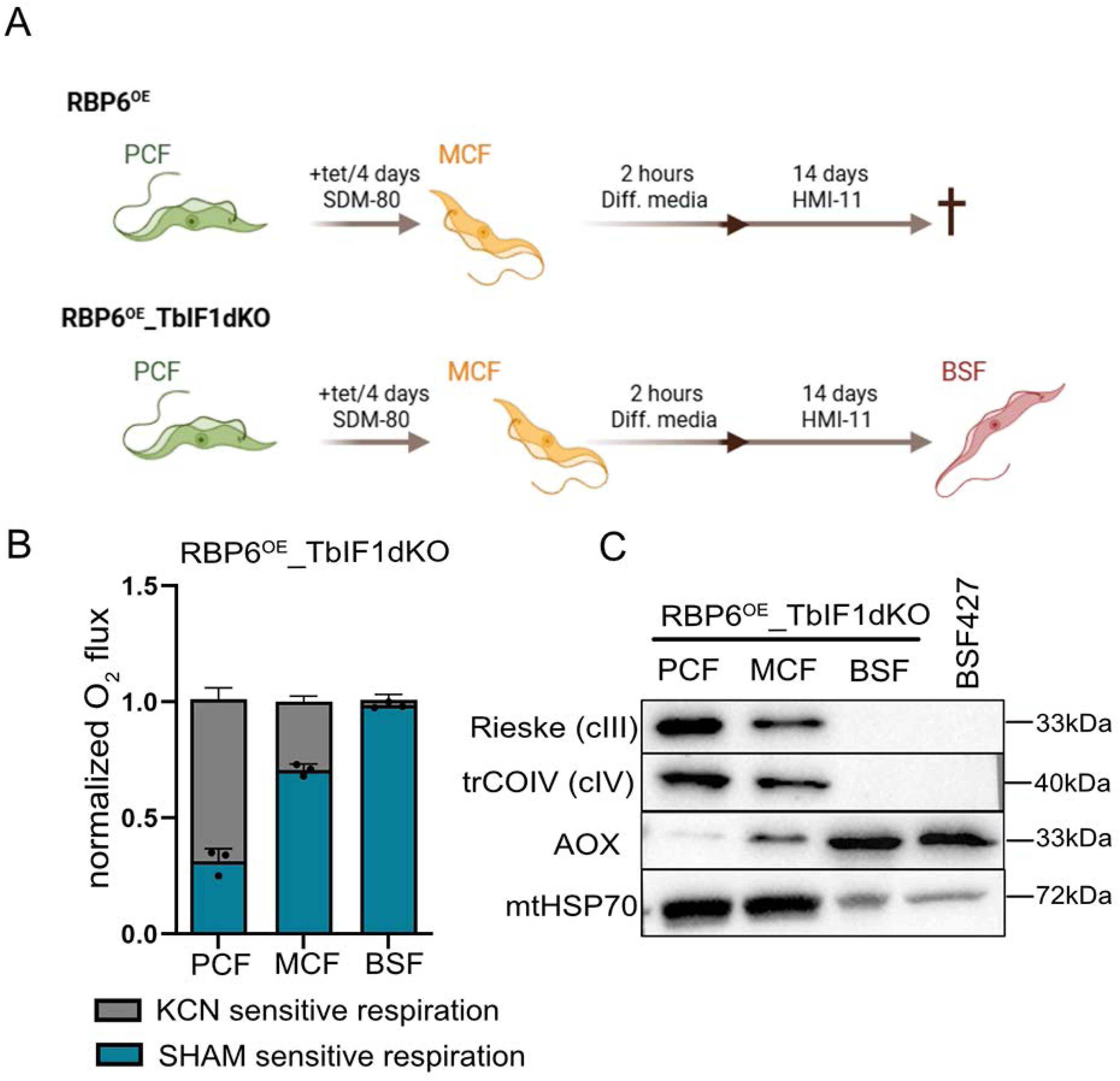
Lack of TbIF1 is an essential prerequisite for successful differentiation to the bloodstream form parasites. (A) Scheme of the differentiation protocol. (B) Oxygen consumption rates in the presence of 10 mM glycerol-3-phopshate in intact of RBP6^OE^_TbIF1 dKO procyclic (PCF), metacyclic (MCF) and bloodstream form (BSF) cells measured using O2k-oxygraph. The ratio of complex IV- and AOX-mediated respiration was determined using KCN, a potent inhibitor of complex IV, and SHAM, a potent inhibitor of AOX. Individual values are shown as dots. (mean ±s.d., n=3) (C) Western blot analysis of whole cell lysates from RBP6^OE^_TbIF1 dKO PCF, MCF, BSF cells and from wild type BSF 427 cells using available antibodies recognizing complex III subunit Rieske, complex IV subunit trCOIV and AOX. Mitochondrial HSP70 (mtHSP70) serves as a loading control between PCF and MCF and BSFs samples.

This study has shown that the regulation of TbIF1 expression and the ATP synthase reversal is a critical factor in physiologically relevant processes such as parasite differentiation. The uninhibited hydrolysis of ATP by the ATP synthase complex plays a key role in maintaining the ΔΨm, particularly in the presence of AOX, which by default causes local membrane depolarization. In addition, the parasite’s differentiation is accompanied by a number of changes in various cellular activities, such as respiration, mROS production, the ADP/ATP ratio, cellular ROS levels, and the activation of AMPK (Figure 7).

**Figure 7.**
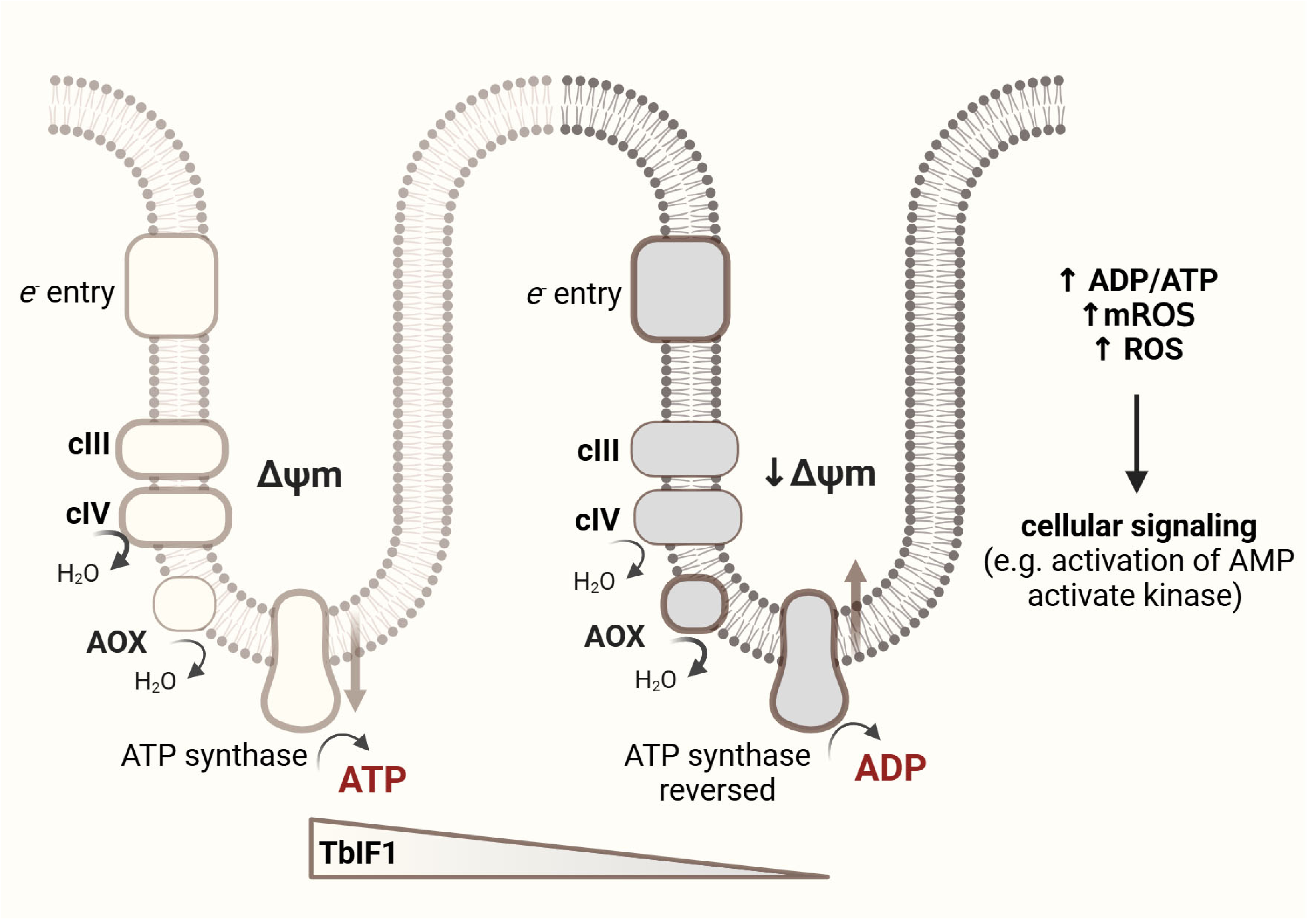
TbIF1 modulates mitochondrial membrane potential and ATP synthase activity in trypanosomes. Schematic representation of mitochondrial electron transport chain (ETC) function and ATP synthase activity under decreasing levels of TbIF1 expression and increasing steady-state levels of alternative oxidase (AOX) during *Trypanosoma brucei* differentiation. Left: In the microenvironment of the mitochondrial cristae, electrons enter the ETC via complex I, alternative NADH dehydrogenases, or complex II, and flow through complexes III (cIII) and IV (cIV). The resulting mitochondrial membrane potential (Δψm) drives ATP synthesis via ATP synthase. Right: During *T. brucei* differentiation, as TbIF1 levels decrease and AOX expression increases, Δψm is reduced. This leads to reversal of ATP synthase activity, causing ATP hydrolysis to maintain Δψm. The shift in bioenergetics results in an increased ADP/ATP ratio, elevated mitochondrial ROS (mROS), and cytosolic ROS, which together contribute to the activation of cellular signaling pathways such as AMP-activated protein kinase (AMPK).

## Discussion

Since its discovery in 1963 [7], IF1 has been extensively studied in various model organisms as a reversible, non-competitive, and unidirectional inhibitor of ATP hydrolysis by ATP synthase [16, 71]. Early studies demonstrated that this protein primarily plays a regulatory role by fine-tuning ATP synthase function, preventing ATP hydrolysis when ΔΨm is compromised due to severe or complete oxygen deprivation or mitochondrial dysfunction associated with a damaged ETC [20, 24]. Considering IF1’s role in pathophysiology, high IF1 expression has also been reported in various tumor cells, where it exerts pro-oncogenic effects and protects cancer cells from hypoxia-induced stress and apoptosis [17, 28]. Recent studies, however, suggest that IF1 also plays a role under physiological conditions, for example, by contributing to metabolic rewiring in T cells [72] or participating in thermogenesis in brown adipose tissue [73]. Here, we investigated the role of IF1 under physiological conditions during the cellular differentiation of *Trypanosoma brucei*. The programmed differentiation of this parasite involves a reversal of ATP synthase activity: insect-stage forms rely on the forward mode of ATP synthase, whereas mammalian-infectious forms depend on the reverse mode of this molecular nanomachine [59]. However, the molecular mechanisms governing this switch remain unknown.

We utilized the ability to induce *T. brucei* differentiation *in vitro* through the overexpression of the RBP6 protein, a process accompanied by significant remodeling of the OXPHOS machinery, including the downregulation of TbIF1 [54, 55]. This observation suggests that modulation of TbIF1 expression plays a crucial role in ensuring proper differentiation. Through the implementation of loss- and gain-of-function experiments, we demonstrated that TbIF1 downregulation is indispensable for parasite metacyclogenesis, while its sustained expression impairs the parasités ability to differentiate.

A hallmark of RBP6-induced mitochondrial metabolic remodeling is the early upregulation of AOX following RBP6 induction. Due to its high maximal velocity (Vmax) for ubiquinol [74], *T. brucei* AOX alters electron flow within the ETC. This contrasts with the *Ciona intestinalis* AOX, which is commonly used in many human disease models [75], has a low Vmax [76], and typically becomes active only under conditions of ETC impairment [75]. Upon the RBP6 induction and in agreement with the increased expression of AOX, NADH-linked substrates induced a lower degree of inner mitochondrial membrane polarization compared to non-differentiating cells. This observation suggests a partial redirection of electrons toward AOX, an enzyme that does not contribute to pmf. Consequently, AOX activity may cause localized depolarization, potentially leading to a reversal of ATP synthase activity within discrete mitochondrial microdomains, despite the overall retention of OXPHOS capacity at the organellar level. Emerging evidence supports the notion that ATP hydrolysis and synthesis can coexist within individual mitochondria, indicating the potential for spatially segregated zones of opposing ATP synthase activity within the same mitochondrial network [29–32]. The extent of the ATP synthase reversal depends on the levels of IF1. In *T. brucei* during the RBP6-induced differentiation, the expression of TbIF1 is strongly downregulated allowing ATP synthase to reverse when ΔΨm is decreased as electrons are partially redirected to non-proton pumping AOX. In addition to the decreased levels of ΔΨm, ATP required for the reversed ATP synthase is produced by mitochondrial substrate-level phosphorylation pathway [50]. During RBP6-induced differentiation, proline oxidation is markedly upregulated, leading to the production of α-ketoglutarate, which serves as a substrate for succinyl-CoA synthetase, an enzyme that generates ATP. This proline-driven respiration is further enhanced in the RBP6^OE^_TbIF1dKO cell line, where TbIF1 is entirely absent. The importance of ATP synthase reversal during the parasite’s differentiation is evident from TbIF1 overexpression experiments, in which RBP6OE_TbIF1OE cells exhibited significantly reduced ΔΨm, suggesting that IF1 overexpression impairs the mitochondria’s ability to maintain ΔΨm via reverse ATP synthase activity during differentiation. Conversely, during the RBP6-induced differentiation, the TbIF1 expression downregulation facilitates ATP synthase reversal, enabling the energetic adaptation required for the metacyclogenesis. The interplay between AOX and IF1 levels allowing ATP synthase reversal might be an overlooking phenomenon in studies of various disease etiologies. Xenotopic expression of AOX has been employed as a tool to investigate mitochondrial dysfunction across various disease models, demonstrating notable rescue effects in some cases, while exacerbating the condition in others (reviewed in [75]). Given that IF1 expression varies considerably among different cell lines, cell types, and tissues (reviewed in [16]), the potential for ATP synthase reversal could be considered when examining disease etiologies.

IF1 has traditionally been considered a unidirectional inhibitor of ATP synthase. However, this view has been challenged, as some recent reports have shown that human IF1 might be capable of inhibiting ATP synthase activity *in vivo*, with this function being regulated by the phosphorylation of serine at the position 14 of the mature human IF1 [15, 71, 77]. Interestingly, this regulatory serine is conserved in humans and mice but is not universally present across the mammalian clade, casting doubt on the generality of this regulatory mechanism [9]. Notably, this serine residue is also absent in *Trypanosoma brucei*. Consistent with this, our data do not support the hypothesis that TbIF1 also inhibits the forward (synthase) mode of ATP synthase. In *T. brucei* procyclic forms, overexpression of TbIF1 had no effect on ATP production via OXPHOS, nor did it increase the ΔΨm [45]. Similarly, in *Toxoplasma gondii*, overexpression of IF1 did not alter ATP synthase activity, ADP/ATP ratio, or ΔΨm [78]. These findings suggest that if human IF1 functions also as an inhibitor of the forward mode of ATP synthase, this role may have evolved specifically in certain multicellular organisms, potentially as a part of a more complex regulatory network governing mitohormesis.

Upon mitochondrial depolarization, reversed ATP synthase consumes ATP, initially supplied by the mitochondrial substrate-level phosphorylation pathway. However, if the mitochondrial membrane potential (ΔΨm) falls below a critical threshold, the ATP/ADP carrier also reverses, importing ATP into the mitochondrial matrix [6, 79, 80]. Therefore, it is expected that the F_o_F_1_- ATPase activity influences the cellular ADP/ATP ratio, a critical cellular indicator of the cell’s energetic status. Indeed, as early as the first day following RBP6 induction, an increase in the ADP/ATP ratio was observed in both the parental RBP6^OE^ cell line as well as in the RBP6^OE^_TbIF1dKO cells. In general, cells are highly sensitive to fluctuations in the ADP/ATP ratio, as an increase in this ratio signals energetic stress. Elevated ADP/ATP ratios are typically accompanied by increased AMP levels, which are sensed by AMP-activated protein kinase (AMPK), a key regulator of cellular homeostasis [81]. AMPK is responsible for monitoring the energy status of the cell and modulating gene expression to help the cell adapt to reduced ATP levels. During the differentiation of RBP6^OE^ cells, we observed AMPK α1 subunit phosphorylation as an indicator of AMPK activation, with a more pronounced response in the RBP6^OE^_TbIF1dKO line, whereas the activation was not detected in the RBP6^OE^_TbIF1^OE^ line.

While AMPK activation is typically triggered by elevated AMP levels, there is also evidence that increased ROS levels can contribute to its activation [68, 69]. This observation is consistent with the results of our study, which revealed an increase in physiological levels of cellular ROS during differentiation, with higher levels detected in the RBP6^OE^_TbIF1dKO cell line and lower levels in the RBP6^OE^_TbIF1^OE^ cells. *T. brucei* AMPK has been identified as a positive regulator of metacyclogenesis, as RNAi-mediated knockdown of all three AMPK subunits significantly reduced the expression of metacyclic-specific genes [57]. Furthermore, AMPK activation is imperative for the differentiation of the proliferative bloodstream form into the quiescent, cell cycle-arrested stumpy form [67]. These stumpy forms are primed for infection of the insect host and exhibit a distinct gene expression profile suited for transmission. A key shared feature of the insect-stage metacyclic form and the bloodstream stumpy forms is that both are non-dividing, cell cycle arrested, exhibit reduced metabolism, and possess a transcriptome prepared for transition to a new host. This suggests that the signaling pathways driving cellular quiescence may be comparable between the differentiation of insect and mammalian forms of the parasite. The involvement of AMPK in parasite differentiation underscores a remarkable evolutionary adaptation, wherein canonical stress-response pathways are repurposed to drive developmental transitions and enhance parasite fitness throughout its complex life cycle.

In addition to its established role as a unidirectional inhibitor, IF1 also plays a crucial structural function in mitochondrial cristae organization [24, 82]. In mammals, mitochondria are characterized by the presence of lamellar cristae, which feature long rows of ATP synthase dimers arranged along their edges [83]. This configuration generates pronounced membrane curvature that promotes the formation of a cristae environment optimized for OXPHOS. Mammalian ATP synthase typically forms a type I dimer with an angle of approx. 86° between its two central stalks [84, 85]. The dimer is characterized by the position of the peripheral stalks, which extends along the longitudinal axis of the dimer. A noteworthy finding is that dimeric IF1, in its active form, spans the interface between adjacent F1 domains of neighboring dimers, forming an inter-dimeric bridge. These tetrameric assemblies have been visualized using cryo-electron microscopy [21, 85]. Although the precise biological significance of such inhibited oligomeric complexes remains uncertain, it is plausible that IF1 contributes to their stabilization, thereby supporting cristae integrity. Consistent with this hypothesis, cells lacking IF1 have been reported to exhibit altered cristae morphology [24, 26, 27, 86].

In *Trypanosoma* species, ATP synthase adopts a distinct type IV dimer configuration. This form is characterized by an angle of approximately 60° between the two monomers and the lateral displacement of peripheral stalks to opposite sides of the dimer plane [87]. As a result, dimers positioned at the edges of the discoidal cristae are inclined at a 45° angle relative to the row axis, forming short helical rows composed of approximately three to six dimers [88]. Structural data from *Euglena* indicate that, unlike the mammalian IF1 which bridges adjacent dimers, the IF1 dimer in these organisms binds individual monomers within the same dimer forming intra-dimeric bridge [89]. The actual dimeric interface between two ATP synthase monomers comprises the subunit e/g module, which is stabilized by bound cardiolipin molecules [87]. The dimer stability seems not to be affected by the absence of TbIF1 as we did not observe changes in ATP synthase oligomerization using BN-PAGE in cells lacking TbIF1. Neither electron microscopy analysis revealed substantial alterations in cristae structure in RBP6^OE^_TbIF1dKO. However, further detailed studies are warranted to assess subtle architectural changes. In *Toxoplasma gondii*, ATP synthase assembles into yet another distinct form, type III dimers, with laterally offset peripheral stalks [90]. In this case, the monomer-monomer angle is approximately 19°, which contrasts with the wider angle observed in type I or IV dimers. Structural data show that dimeric IF1 binds to both monomers within the same dimer, forming an intra-dimeric bridge. Additionally, ATP synthase dimers in *T. gondii* organize into cyclic trimers of dimers. Notably, the dimer-dimer interface involves contacts within the luminal regions and does not incorporate IF1. Nevertheless, knock-out of IF1 in *T. gondii* tachyzoites led to a modest reduction in cristae density, suggesting a potential role for IF1 in maintaining cristae structure [78].

In summary, our study demonstrates that the role of IF1 in controlling ATP synthase activity is more significant in regulating cellular energy metabolism under physiological conditions than previously anticipated. In *T. brucei*, the programmed regulation of TbIF1 expression is a critical prerequisite for successful progression through the organism’s life cycle. Our data strongly emphasize the pivotal role of the ATP synthase/IF1 axis in cellular signaling.

## Acknowledgment

We would like to thank Martina Slapničková for excellent technical support and prof. Christos Chinopoulos (Semmelweis University, Budapest) for stimulating discussions. We would also like to express our gratitude to the Biology Centre core facilities, namely to the Laboratory of Electron Microscopy, to the Laboratory of Microscopy and Histology and to the Laboratory of Analytical Biochemistry and Metabolomics.

## Supplementary Figures

**S1.**
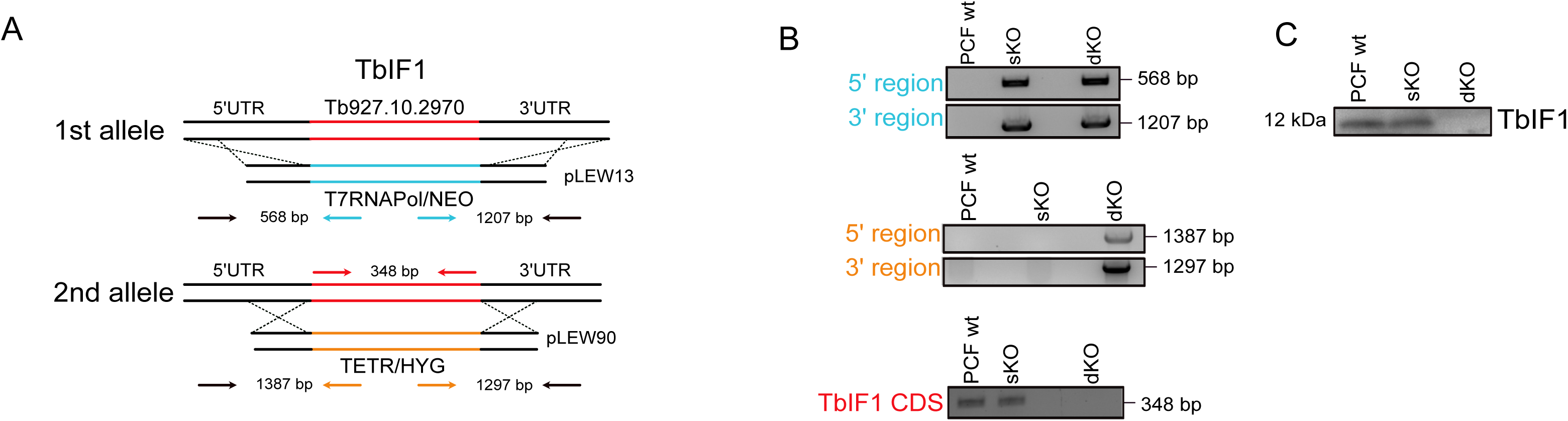
Generation of TbIF1 double knock-out (dKO). (A) Scheme of the dKO generation followed by incorporation of RBP6^OE^ cassette to the rRNA spacer. (B) PCR validation of the correct TbIF1 gene allele replacement with pLEW13 and pLEW90 cassettes. (C) Western blot analysis using anti-TbIF1 specific antibody validating the absence of TbIF1 protein in the RBP6OE_TbIF1dKO cells.

**S2.**
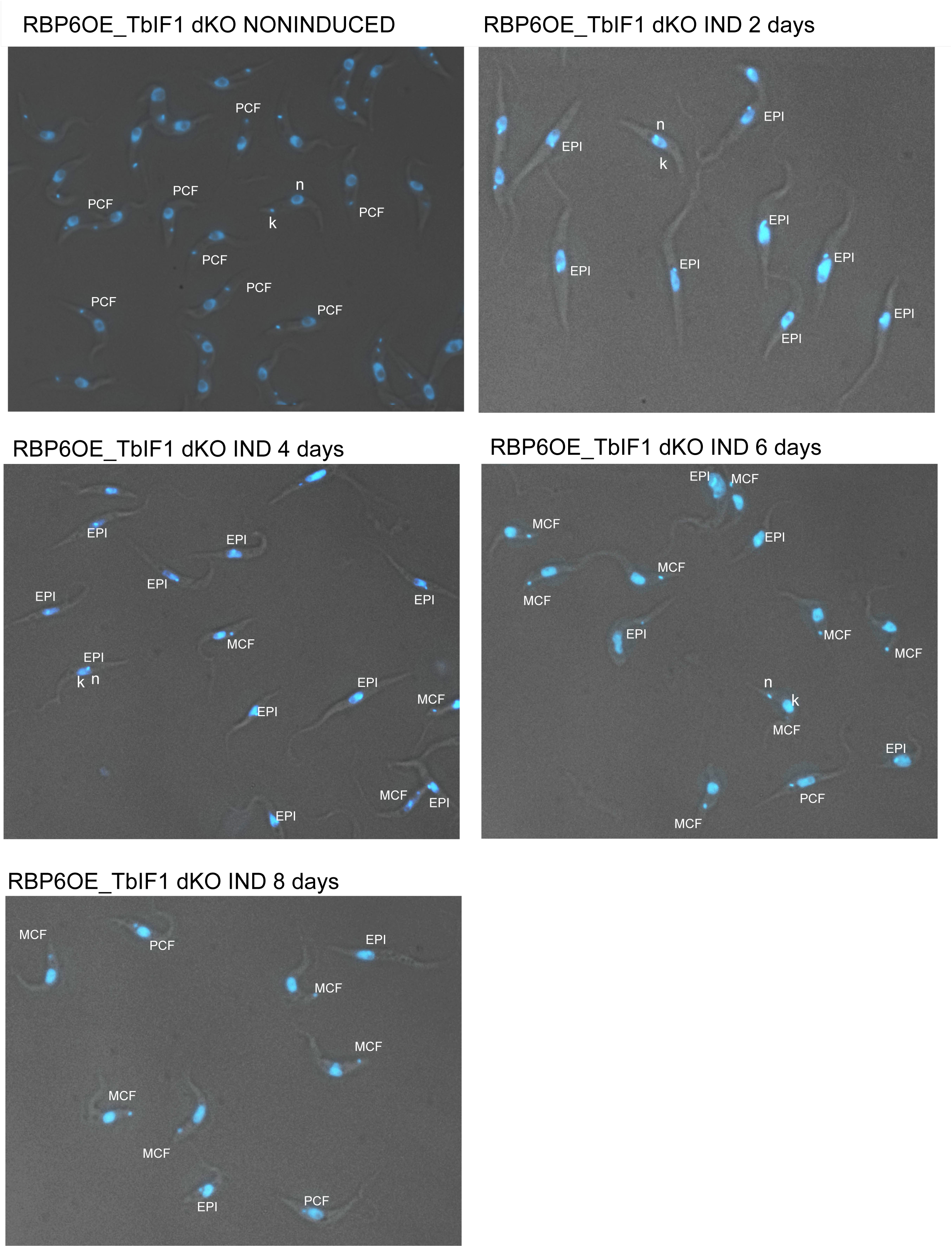
Morphological progression of *T. brucei* induced by RBP6 overexpression. Morphological stages of *T. brucei* during RBP6 overexpression. Representative images show differentiation from procyclic forms (PCF) to epimastigote (EPI) and metacyclic forms (MCF) over an 8-day induction period. Samples include non-induced and induced RBP6^OE^_TbIF1 dKO parasites at 2, 4, 6, and 8 days. Panels are labeled to indicate the observed morphological forms

**S3.**
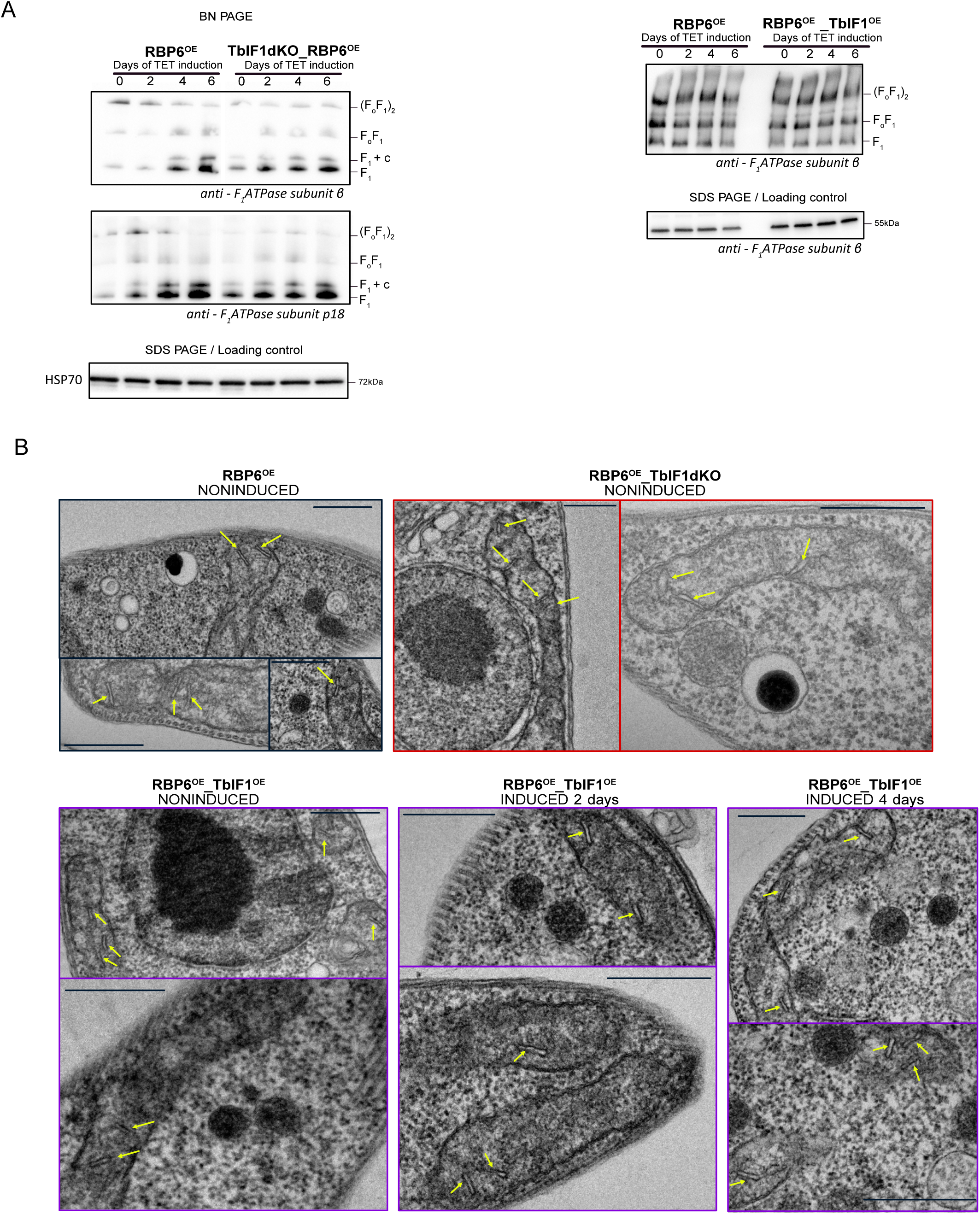
Modulation of TbIF1 expression has no effect on the steady state of the ATP synthase monomer and dimer. (A) A western blot analysis of mitochondrial lysates resolved under native conditions using anti-subunit β antibody recognizing the F1 moiety, as well as the monomeric and dimeric ATP synthase complex. Mitochondrial lysates were also analyzed by SDS PAGE and western blot analysis using mtHSP70 or anti-subunit β antibody as a loading control. (B) Representative transmission electron micrographs of sections of RBP6^OE^ and RBP6^OE^_TbIF1dKO noninduced cells as well as RBP6^OE^_TbIF1^OE^ cells induced for 0, 2 and 4 days. Mitochondrial cristae are marked with yellow arrows. Scale bar 500 nm.

